# Non-invasive, label-free optical analysis to detect aneuploidy within the inner cell mass of the preimplantation embryo

**DOI:** 10.1101/2020.11.01.364133

**Authors:** Tiffany C. Y. Tan, Saabah B. Mahbub, Carl A. Campugan, Jared M. Campbell, Abbas Habibalahi, Darren J. X. Chow, Sanam Mustafa, Ewa M. Goldys, Kylie R. Dunning

## Abstract

**Study question:** Can label-free, non-invasive optical imaging by hyperspectral microscopy discern between euploid and aneuploid cells within the inner cell mass of the mouse preimplantation embryo?

**Summary answer:** Hyperspectral microscopy shows a variance in metabolic activity which enables discrimination between euploid and aneuploid cells.

**What is known already:** Euploid/aneuploid mosaicism affects up to 17.3% of human blastocyst embryos with trophectoderm biopsy or spent media currently utilised to diagnose aneuploidy and mosaicism in clinical in vitro fertilisation. Based on their design, these approaches will fail to diagnose the presence or proportion of aneuploid cells within the fetal lineage (inner cell mass (ICM)) of some blastocyst embryos.

**Study design, size, duration:** The impact of aneuploidy on cellular metabolism of primary human fibroblast cells and mouse embryos was assessed by a fluorescence microscope adapted for imaging with multiple spectral channels (hyperspectral imaging). Primary human fibroblast cells with known ploidy were subjected to hyperspectral imaging to record native cell fluorescence (euploid n= 467; aneuploid n= 969). For mouse embryos, 50-70 individual euploid and aneuploid blastomeres (8-cell stage embryo) and chimeric blastocysts (40-50 per group: euploid; aneuploid; or 1:1 and 1:3 ratio of euploid:aneuploid) were utilised for hyperspectral imaging.

**Participants/materials, setting, methods:** Two models were employed: (i) Primary human fibroblasts with known karyotype and (ii) a mouse model of embryo aneuploidy where mouse embryos were treated with reversine, a reversible spindle assembly checkpoint inhibitor, during the 4-to 8-cell division. Individual blastomeres were dissociated from reversine treated (aneuploid) and control (euploid) 8-cell embryos and either imaged directly or used to generate chimeric blastocysts with differing ratios of euploid:aneuploid cells. Individual blastomeres and embryos were subjected to hyperspectral imaging. Changes in cellular metabolism were determined by quantification of metabolic cofactors (inferred from their autofluorescence signature): reduced nicotinamide adenine dinucleotide (NAD(P)H), flavins with the subsequent calculation of the optical redox ratio (ORR: Flavins/[NAD(P)H + Flavins]). Mathematical algorithms were applied to extract features from the autofluorescence signals of each cell/blastomere/inner cell mass to discriminate between euploid and aneuploid.

**Main results and the role of chance:** An increase in the relative abundance of NAD(P)H with a decrease in flavins led to a significant reduction in the ORR for aneuploid cells in both primary human fibroblasts and individual mouse blastomeres (P < 0.05). Mathematical algorithms were able to achieve good separation between (i) euploid and aneuploid primary human fibroblast cells, (ii) euploid and aneuploid mouse blastomeres cells and (iii) euploid and aneuploid chimeric blastocysts and (iv) 1:1 and 1:3 chimeric blastocysts. The accuracy of these separations was supported by receiver operating characteristic curves with areas under the curve of 0.85, 0.99, 0.87 and 0.88, respectively. We believe that the role of chance is low as multiple cellular models (human somatic cells and mouse embryos) demonstrated a consistent shift in cellular metabolism in response to aneuploidy as well as the robust capacity of mathematical features to separate euploid and aneuploid cells in a statistically significant manner.

**Limitations, reasons for caution:** There would be added value in determining the degree of embryo mosaicism by sequencing the inner cell mass (ICM) of individual blastocysts to correlate with metabolic profile and level of discrimination achieved using the mathematical features approach.

**Wider implications of the findings:** Hyperspectral imaging was able to discriminate between euploid and aneuploid human fibroblasts and mouse embryos. This may lead to the development of an accurate and non-invasive optical approach to assess mosaicism within the ICM of human embryos in the absence of fluorescent tags.

**Study funding/competing interest(s):** K.R.D. is supported by a Mid-Career Fellowship from the Hospital Research Foundation (C-MCF-58-2019). This study was funded by the Australian Research Council Centre of Excellence for Nanoscale Biophotonics (CEI40100003). The authors declare that there is no conflict of interest.

## 1. Introduction

During the early days of life, errors occur during mitotic cell division resulting in the emergence of aneuploidy: a deviation from the expected 46 chromosomes. Mitotic aneuploidy during preimplantation embryo development can lead to mosaic blastocyst embryos, containing both euploid cells (expected number of chromosomes) and aneuploid cells. The incidence of mosaicism in the blastocyst is as high as 30-40% (Fragouli, *et al.*, 2011, Huang, *et al.*, 2019, Northrop, *et al.*, 2010), with euploid/aneuploid mosaicism reported to vary from to 17.3% (Capalbo, *et al.*, 2014, Fragouli, *et al.*, 2011, Huang, *et al.*, 2019, Johnson, *et al.*, 2010, Northrop, *et al.*, 2010). Animal studies show that while a lower proportion of aneuploid cells in the embryo is tolerated, a higher proportion of such cells leads to an increased incidence of pregnancy loss (Bolton, *et al.*, 2016). Thus, the determination of aneuploid cell proportions would likely aid in stratification of embryo quality and improve clinical in vitro fertilisation (IVF) success. To this end, genetic testing of preimplantation human embryos is widely employed albeit amongst considerable debate over the accuracy and safety of these clinical tools.

Preimplantation genetic testing for aneuploidy (PGT-A) is the most widely used method to detect aneuploidies in clinical IVF. This involves an invasive biopsy of trophectoderm cells followed by sequencing. However due to the random nature of mosaicism, this biopsy will not provide a measure of the proportion of aneuploid cells in the ICM (fetal cells) or the remainder of the TE (placental cells) in some embryos (Gleicher, *et al.*, 2017). This is because aneuploid cells can be restricted to the TE, while the ICM is chromosomally normal (and vice versa), resulting in false positives (or negatives)(Vera-Rodriguez, *et al.*, 2017). This can lead to 1) erroneous disposal of false-positive embryos that would have otherwise resulted in a healthy offspring or 2) transfer of false-negative embryos, which may result in implantation failure, pregnancy loss or foetal abnormality. Further, transfer of biopsied embryos leads to a 3-fold increased risk of preeclampsia: a pregnancy complication with elevated risk of morbidity and mortality for mother and fetus and a life-time increased risk of cardiovascular disease (Mastenbroek, *et al.*, 2011).

More recently, an alternative method for preimplantation genetic testing has emerged using cell-free DNA (cfDNA) released by embryos into the culture media (Rubio, *et al.*, 2019, Vera-Rodriguez, *et al.*, 2018). This approach, termed non-invasive preimplantation genetic testing for aneuploidy (niPGT-A), has shown high concordance with TE biopsy (Feichtinger, *et al.*, 2017, Huang, *et al.*, 2019). However, other publications report highly variable results which is thought to arise from differences in culture method, level of embryo mosaicism (Xu, *et al.*, 2016), and potential contamination of maternal DNA in the spent medium (Vera-Rodriguez, *et al.*, 2018). Importantly, this technique assumes that cfDNA originates equally from the ICM and TE (Kuznyetsov, *et al.*, 2020), which has not yet been proven. Therefore, it is probable that this approach will fail in some embryos to detect aneuploidy within the ICM (false negative) or mis-diagnose genetically normal ICM (false positive) (Gleicher, *et al.*, 2019). Currently, it appears that accurate detection of mosaicism within the divergent lineages of the blastocyst embryo can only be achieved through destructive dissection of the embryo (Gleicher, *et al.*, 2017, Taylor, *et al.*, 2016).

Aneuploidy results in altered cell metabolism in primary human fibroblasts and cancer cells (Newman, *et al.*, 2019, Sheltzer, 2013, Warburg, 1956), yeast (Sheltzer, *et al.*, 2011, Torres, *et al.*, 2007), and human embryos (Fragouli, *et al.*, 2015, Picton, *et al.*, 2010). As metabolites and co-enzymes of metabolic pathways are naturally fluorescent, including nicotinamide adenine dinucleotide (NAD), nicotinamide adenine dinucleotide phosphate (NADPH) and flavins, there is potential to exploit this fluorescence as a non-invasive indicator of aneuploidy. Importantly, these endogenous fluorophores can be directly distinguished by their excitation and emission profiles without the need for exogenous fluorescence labels (Campbell, *et al.*, 2019). Both NAD(P)H and flavins are increasingly used to reflect cellular metabolism in oocytes and early embryos (Aparicio, *et al.*, 2013, Dumollard, 2004, Sugimura, *et al.*, 2014, Sutton-McDowall, *et al.*, 2016, Thompson, *et al.*, 2016).

Hyperspectral microscopy is a label-free approach used to non-invasively identify native endogenous fluorophores within a cell (Mahbub, *et al.*, 2017). This technique excites endogenous fluorophores with a wide range of excitation and emission wavelengths as opposed to using single (or dual) excitation wavelength(s). Notably, hyperspectral imaging has demonstrated the capability to discriminate between different types of cancer cells (Campbell, *et al.*, 2019, Gosnell, *et al.*, 2016b), bovine embryos with good and poor developmental potential (Santos Monteiro, *et al.*, 2020, Sutton-McDowall, *et al.*, 2017), as well as quantifying the metabolic content of oocytes from young and aged mice (Bertoldo, *et al.*, 2020).

In the present study, we determine whether hyperspectral imaging is able to discriminate between euploid and aneuploid primary human fibroblasts and mouse embryos. Aneuploid mouse embryos were generated using reversine, a spindle assembly inhibitor, and used to create mosaic blastocysts with different ratios of euploid/aneuploid cells (Bolton, *et al.*, 2016). As current clinical tools are limited in their capacity to detect aneuploidy within the foetal cell lineage of the blastocyst, the aim of this study was to determine whether hyperspectral imaging coupled with mathematical feature extractions was able to discriminate between the ICM of embryos with varying proportions of aneuploid cells. Further, we assessed the safety of this approach using post-transfer viability.

## 2. Materials and methods

All reagents were purchased from Sigma-Aldrich (St. Louis, MO, USA) unless stated otherwise.

### 2.1 Animals and ethics

Female (21-23 days) and male (6-8 weeks old) CBA x C57BL/6 first filial (F1) generation (CBAF1) mice were obtained from Laboratory Animal Services (LAS; University of Adelaide, SA, Australia) and maintained on a 12 h light:12 h dark cycle with rodent chow and water provided *ad libitum*. All experiments were approved by the University of Adelaide’s Animal Ethics Committee and were conducted in accordance with the Australian Code of Practice for the Care and Use of Animals for Scientific Purposes.

### 2.2 Media for gamete and embryo handling and culture

All gamete and embryo culture took place in media under paraffin oil at 37 °C in a humidified atmosphere of 5 % O_2_, 6 % CO_2_ with the balance as N_2_. All media were pre-equilibrated at least 4 h prior to use. All handling procedures were carried out at 37 °C. Mouse reproductive tissues were collected in Research Wash Medium (ART Lab Solutions, SA, Australia) supplemented with 4 mg/ml of low fatty acid bovine albumin (BSA; MpBio). Research Fertilisation Medium and Cleave Medium (ART Lab Solutions, SA, Australia) were supplemented with 4 mg/ml low fatty acid BSA and used for *in vitro* fertilisation (IVF) and embryo culture, respectively. Embryo vitrification and warming were carried out in Handling Medium (HM; alpha Minimal Essential Medium; αMEM; Gibco by Life Technologies, CA, USA) containing: 10 mM HEPES-buffered αMEM medium supplemented with 6 mM NaHCO3, 50 mg/L gentamicin sulfate, 5.56 mM glucose and 2 mM glutamax. Before use, HM was supplemented with 5 mg/ml low fatty acid BSA.

### 2.3 Isolation of COCs and in vitro fertilisation

Female mice were administered intraperitoneally (i.p.) with 5 IU equine chorionic gonadotrophin (eCG; Braeside, VIC, Australia), followed by 5 IU (I.P.) human chorionic gonadotrophin (hCG; Kilsyth, VIC, Australia) 48 h later. Mice were culled by cervical dislocation 14 h post-hCG administration and oviducts removed. Ovulated COCs were harvested by gently puncturing the ampulla using a 29-gauge needle. Male mice with proven fertility were culled by cervical dislocation one hour prior to IVF. Spermatozoa were released from the vas deferens and the caudal region of the epididymis by blunt dissection in Research Fertilisation Medium and allowed to capacitate for 1 h at 37 °C, 5 % O_2_, 6 % CO_2_ with the balance as N_2_. Mature COCs were then co-cultured with 10 μl of capacitated spermatozoa for 4 h at 37 °C in an atmosphere of 5 % O_2_, 6 % CO_2_ with the balance as N_2_. Resulting presumptive zygotes were transferred into Research Cleave Medium (in groups of 10; 2 μl per embryo) and cultured at 37 °C in an atmosphere of 5 % O_2_, 6 % CO_2_ with the balance as N_2_. Fertilisation rate was scored 24 h later, with embryos allowed to develop to the 4-cell or blastocyst stage.

### 2.4 Euploid/ Aneuploid/ Chimeric Embryo Generation

Generation of chimeric embryos was performed as described in (Bolton, *et al.*, 2016), using reversine, a biomolecule inhibitor of monopolar spindle 1-like 1 kinase, which is crucial for the normal functioning of the spindle assembly checkpoint (SAC) (Santaguida, *et al.*, 2010). Briefly, during the 4-to 8-cell division, embryos were cultured in the absence (euploid) or presence of 0.5 μM reversine (aneuploid) diluted in Research Cleave Medium. A group of non-treated 8-cell embryos were set aside to culture directly to the blastocyst stage to investigate whether the presence of the zona pellucida altered the autofluorescence profile of embryos. The remaining 8-cell embryos were incubated in 0.05 % pronase diluted in MOPS to remove the zona pellucida. Individual blastomeres were then separated in calcium and magnesium free medium (Research Wash Medium without calcium and magnesium) using a STRIPPER Micropipette Handle (CooperSurgical, Trumbull, Connecticut) fitted with a 30-35 μm biopsy pipette (TPC micropipettes; CooperSurgical, Trumbull, Connecticut). Following separation, individual blastomeres were washed thoroughly in Research Wash Medium and either subjected to imaging using the hyperspectral microscope or reaggregated in 2% phytohaemagglutinin M (ThermoFisher Scientific, Waltham, Massachusetts) in Research Cleave Medium to generate chimeric blastocysts: euploid; aneuploid; 1:1 or 1:3 ratio of euploid:aneuploid. Agglutinated chimeric 8-cell embryos were then cultured overnight at 37 °C, 5 % O_2_, 6 % CO_2_ with the balance as N_2_. Chimeric embryos were only included if there was aggregation of all 8 blastomeres and development to the morula stage. This is to ensure imaging was only performed on blastocysts that resulted from the agglutination of all 8 cells, and thus reflective of their allocated ratio of euploid:aneuploid cells.

For hyperspectral imaging of individual blastomeres, 8-cell embryos were vitrified and shipped from the University of Adelaide (UoA) to the University of New South Wales (UNSW) in a dry shipper to maintain cryogenic temperature. For the imaging of individual blastomeres, 8-cell embryos were warmed and individual blastomeres separated using the biopsy micropipette as described above. For hyperspectral imaging of chimeric embryos, chimeric morulas were created at UoA as described above, subjected to vitrification, and shipped in a dry shipper to UNSW. Chimeric morulas were then subjected to embryo warming and allowed to develop to the blastocysts stage prior to hyperspectral imaging.

### 2.5 Embryo Vitrification and Warming

We utilised the CryoLogic vitrification method (CVM) for the vitrification of 8-cell and morula embryos. A NUNC four-well dish (ThermoFisher Scientific, Waltham, Massachusetts) was set up with 600 μl of HM, equilibration solution (ES; HM supplemented with 10 % each ethylene glycol (EG) and DMSO), and vitrification solution (VS; HM supplemented with 16 % each for EG and DMSO and 0.5 M sucrose). After pre-warming to 37 °C, 8-cell or morula embryos were washed twice in HM, followed by transfer into ES for 3 min. Embryos were then transferred into a VS drop for 30-45 s before loaded onto a Fibreplug (Cryologic,Pty. Ltd, Victoria) and immediately vitrified using the CVM system, followed by storage in a straw in a liquid nitrogen tank.

For embryo warming, HM supplemented with decreasing concentrations of sucrose (0.3M, 0.25M and 0.15M) were prewarmed to 37 °C. Storage straws containing 8-cell or morula embryos were kept immersed in liquid nitrogen prior to use. The Fibreplug was removed from the straw and quickly submerged in HM containing 0.3M sucrose for 30 s, followed by transfer to decreasing concentrations of sucrose. Prior to imaging, 8-cell embryos were recovered in Research Cleave Medium for 2 h then dissociated into individual blastomeres as described above. Chimeric embryos at the morula stage were transferred to Research Cleave Medium and cultured overnight at 37 °C, 5 % O_2_, 6 % CO_2_ with the balance as N_2_ until they reached the blastocysts stage. Survival of the warmed embryos was assessed morphologically, based on the presence of inner cell mass and blastocoel cavity. Blastocysts were then subjected to hyperspectral imaging.

### 2.6 Tissue Culture of Primary Human Fibroblasts

Primary human fibroblasts (Coriell Institute for Medical Research, USA) with known karyotypes (46, XY (GMO0970); 46, XX (GMO3525); triploid (GM01672); trisomy: 13 (GM02948), 18 (GM01359), 21 (AG06922), XXX (GM04626), and XXY (GM03184)) were cultured in MEM-E (Minimum Essential Medium Eagle with Earle’s salts, 2.2 g/L sodium bicarbonate and, supplemented with 2 mM L-glutamine (Gibco), 1% MEM Non-Essential Amino Acids Solution (Gibco, ThermoFisher Scientific, Waltham, Massachusetts), 100 U/ml penicillin-streptomycin, and 10% fetal calf serum (FCS; Gibco). Cell cultures were maintained in 5 % CO_2_ in air at 37°C. Using a 35 mm glass bottom dish (Ibidi, Martinsried, Planegg, Germany), 1 × 10^5^ cells were plated and cultured overnight. Prior to imaging, cells were washed twice with PBS without calcium and magnesium. Cells were imaged in Hank’s balanced salt solution (Gibco, ThermoFisher Scientific, Waltham, Massachusetts).

### 2.7 Hyperspectral and differential interference contrast (DIC) Imaging

In this work, the hyperspectral microscopy system was built by adapting a standard fluorescence microscope (Olympus iX83™) fitted with a 60x silicon U12™ series objective to capture and characterised fluorescence of native endogenous fluorophores found within cells (Gosnell, *et al.*, 2016a, Gosnell, *et al.*, 2016b). A hyperspectral excitation lamp (Quantative™, Australia) comprising multiple low-powered light-emitting diodes (LEDs) with selected bands of excitation wavelengths produced by the LEDs (centred at 334 nm, 365 nm, 385 nm, 395 nm, 405 nm, 415 nm, 425 nm, 435 nm, 455 nm, 475 nm, and 495 nm, each about 10 mm wide) were used to excite cellular autofluorescence. The subsequent emission, through the use of three epifluorescence filter cubes with the emission bands centred at 447 nm (60 nm wide), 587 nm (35 nm wide), and 700 nm long pass, allowed us to record the autofluorescence signal of cell samples in a number of defined narrowbands (±5 nm) to generate specific and defined spectral channels, covering a broad excitation (345 nm-505 nm) and emission (414 nm-675 nm) wavelength ranges (see Table I for details of spectral channels). In total, 67 specific spectral channels were available to measure single photon-excited emission, or autofluorescence signals of the biological sample. Images were captured by an electron multiplying CCD (Nuvu™ 1024, Canada) operated below −80 °C to minimise sensor-induced noise in the image data. The sensor is made out of 1024×1024 (13 μm × 13 μm) pixels. We used image acquisition times of up to 5 seconds per channel, with multiple averaging (typically 3-5 times) to optimise image quality and minimise any potential photodamage to the cells in each channel. Each data set was supplemented with a brightfield image of the sample which was used as a broad reference.

**Table I.** Specification of spectral channels with their respective excitation, emission, dichroic mirror wavelengths and laser powers for hyperspectral system in UNSW, Sydney, Australia.

| Spectral Channel | Wavelength (nm) | | | Power (μW) | Exposure time (s) |
| --- | --- | --- | --- | --- | --- |
|  | Excitation | Emission | Dichroic mirror |  |  |
| 1 | 345 | 414 | 389 | 2.80 | 0.2 |
| 2 | 345 | 451 | 389 | 2.90 | 0.2 |
| 3 | 345 | 575 | 552 | 2.70 | 1.5 |
| 4 | 345 | 594 | 552 | 2.90 | 2.0 |
| 5 | 345 | 675 | 552 | 2.00 | 2.0 |
| 6 | 490 | 575 | 552 | 3.98 | 0.8 |
| 7 | 505 | 575 | 552 | 8.40 | 0.1 |
| 8 | 490 | 594 | 552 | 4.11 | 0.1 |
| 9 | 505 | 594 | 552 | 8.63 | 0.1 |
| 10 | 490 | 675 | 552 | 3.29 | 0.9 |
| 11 | 505 | 675 | 552 | 6.90 | 2.0 |
| 12 | 358 | 414 | 389 | 2.76 | 2.0 |
| 13 | 371 | 414 | 389 | 3.10 | 2.0 |
| 14 | 377 | 414 | 389 | 2.10 | 2.0 |
| 15 | 381 | 414 | 389 | 0.87 | 2.0 |
| 16 | 358 | 451 | 389 | 2.54 | 2.0 |
| 17 | 371 | 451 | 389 | 3.14 | 2.0 |
| 18 | 377 | 451 | 389 | 2.17 | 2.0 |
| 19 | 381 | 451 | 389 | 0.90 | 2.0 |
| 20 | 358 | 575 | 552 | 3.02 | 2.0 |
| 21 | 371 | 575 | 552 | 4.60 | 2.0 |
| 22 | 377 | 575 | 552 | 8.52 | 2.0 |
| 23 | 381 | 575 | 552 | 10.19 | 2.0 |
| 24 | 391 | 575 | 552 | 6.92 | 2.0 |
| 25 | 397 | 575 | 552 | 8.45 | 0.01 |
| 26 | 400 | 575 | 552 | 9.20 | 2.0 |
| 27 | 403 | 575 | 552 | 4.90 | 2.0 |
| 28 | 406 | 575 | 552 | 9.73 | 2.0 |
| 29 | 412 | 575 | 552 | 13.51 | 2.0 |
| 30 | 418 | 575 | 552 | 14.80 | 2.0 |
| 31 | 430 | 575 | 552 | 10.21 | 2.0 |
| 32 | 437 | 575 | 552 | 14.31 | 2.0 |
| 33 | 457 | 575 | 552 | 8.33 | 2.0 |
| 34 | 469 | 575 | 552 | 11.89 | 2.0 |
| 35 | 476 | 575 | 552 | 12.86 | 2.0 |
| 36 | 358 | 594 | 552 | 3.13 | 2.0 |
| 37 | 371 | 594 | 552 | 4.84 | 2.0 |
| 38 | 377 | 594 | 552 | 8.95 | 2.0 |
| 39 | 381 | 594 | 552 | 10.73 | 2.0 |
| 40 | 391 | 594 | 552 | 7.26 | 2.0 |
|  | Excitation | Emission | Dichroic mirror |  |  |
| 41 | 397 | 594 | 552 | 8.81 | 2.0 |
| 42 | 400 | 594 | 552 | 9.55 | 2.0 |
| 43 | 403 | 594 | 552 | 5.30 | 2.0 |
| 44 | 406 | 594 | 552 | 10.15 | 0.01 |
| 45 | 412 | 594 | 552 | 14.25 | 2.0 |
| 46 | 418 | 594 | 552 | 15.49 | 2.0 |
| 47 | 430 | 594 | 552 | 10.74 | 2.0 |
| 48 | 437 | 594 | 552 | 14.88 | 2.0 |
| 49 | 457 | 594 | 552 | 8.62 | 2.0 |
| 50 | 469 | 594 | 552 | 12.34 | 2.0 |
| 51 | 476 | 594 | 552 | 13.35 | 2.0 |
| 52 | 358 | 675 | 552 | 2.47 | 2.0 |
| 53 | 371 | 675 | 552 | 3.89 | 2.0 |
| 54 | 377 | 675 | 552 | 7.15 | 2.0 |
| 55 | 381 | 675 | 552 | 8.62 | 2.0 |
| 56 | 391 | 675 | 552 | 5.82 | 2.0 |
| 57 | 397 | 675 | 552 | 7.19 | 2.0 |
| 58 | 400 | 675 | 552 | 7.60 | 2.0 |
| 59 | 403 | 675 | 552 | 4.13 | 2.0 |
| 60 | 406 | 675 | 552 | 8.06 | 2.0 |
| 61 | 412 | 675 | 552 | 11.44 | 2.0 |
| 62 | 418 | 675 | 552 | 12.41 | 2.0 |
| 63 | 430 | 675 | 552 | 8.49 | 0.01 |
| 64 | 437 | 675 | 552 | 11.96 | 2.0 |
| 65 | 457 | 675 | 552 | 6.85 | 2.0 |
| 66 | 469 | 675 | 552 | 10.00 | 2.0 |
| 67 | 476 | 675 | 552 | 10.82 | 2.0 |

In this study, we also utilised an additional hyperspectral microscopy system (UoA) to investigate light-induced damage (photodamage) on embryo development. This hyperspectral system was built with a similar concept by adapting to a standard epifluorescence microscope (Nikon Eclipse TiE) and fitted with a multi-LED light source (Prizmatix Ltd, Givat-Shmuel, Israel). These low power LEDs provide 40 spectral channels: 21 excitation wavelength ranges and three emission wavelength filters, covering excitation wavelengths from 340 to 664 nm and emission wavelengths from 440 – 715 nm (see Table II for details of spectral channels). Images were captured by a digital camera C1140, OCRA Flash 4.0 (Hamamatsu) using all 40 spectral channels.

**Table II.**
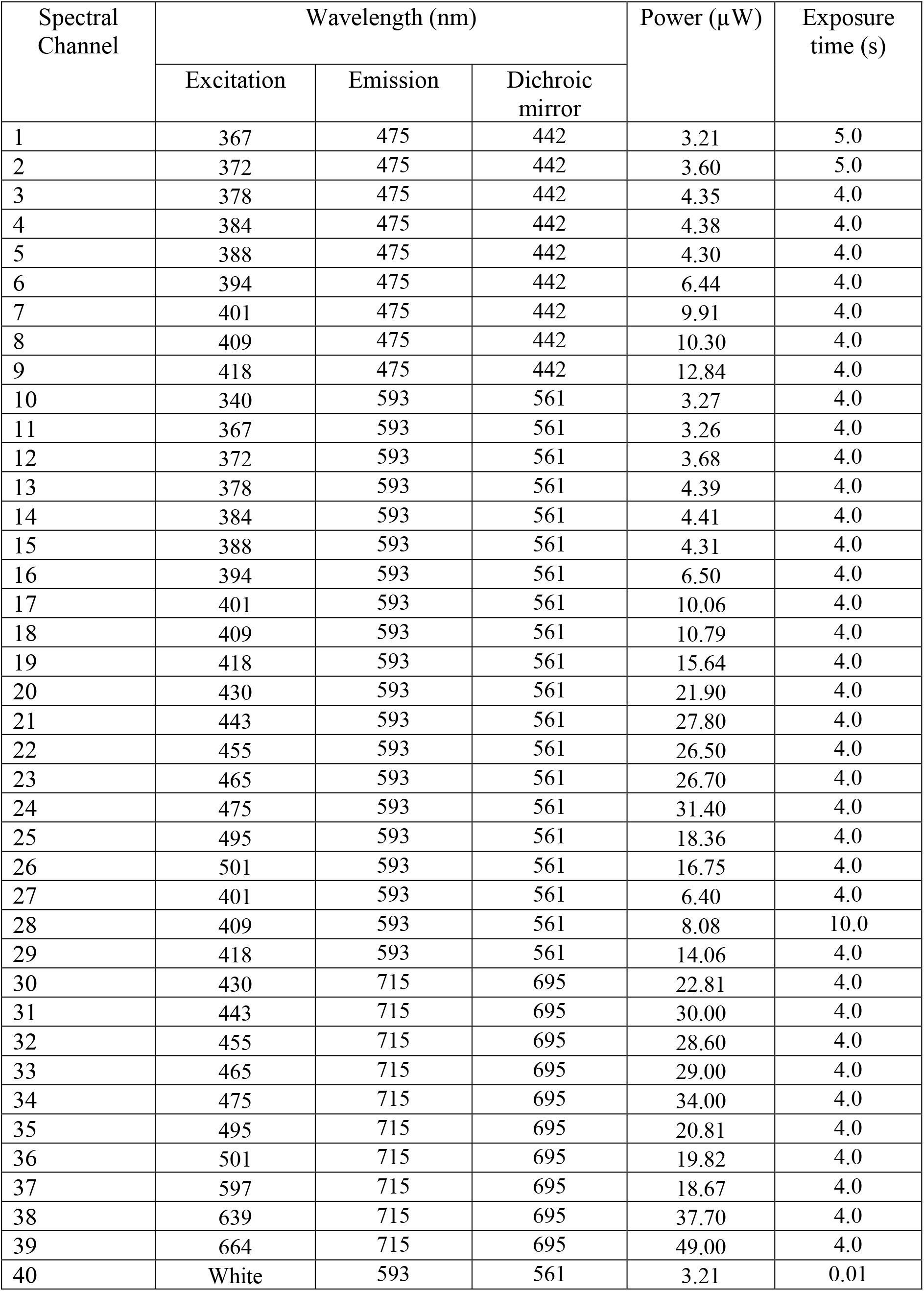
Specification of spectral channels with their respective excitation, emission, dichroic mirror wavelengths and laser powers for hyperspectral system in UoA, SA, Australia.

For imaging, blastomeres, morula embryos and chimeric blastocysts were transferred onto a glass bottomed confocal dish (Ibidi, Martinsried, Planegg, Germany) containing prewarmed Research Wash Medium overlayed with paraffin oil. Hyperspectral images were taken by adjusting the input light beam to specifically focus on the equatorial plane of blastomeres and morula embryos (i.e. widest diameter) and the ICM of individual blastocysts.

### 2.8 Hyperspectral Data Analysis

The analysis of hyperspectral image data in this study included image preparation, classification and unmixing of individual fluorophores. Firstly, image preparation was carried out to remove image artefacts, such as background fluorescence, Poisson’s noise, dead or saturated pixels, and illumination curvature across the field of view, as described in detail in previous work (Habibalahi, *et al.*, 2019, Mahbub, *et al.*, 2017, Rehman, *et al.*, 2017). At the beginning of each experiment, two calibration images were captured using the hyperspectral system: a “background” reference image of a culture dish with medium only, and another with calibration fluid only. The “background” reference image was included and subtracted from all images with cells to remove any potential, unavoidable background signals generated from the glass-bottomed dish or potential fluorescence contamination of microscope optics, which may form an additional fluorescence source to all spectral images. The microscope system was calibrated with a mixture of 30 μM NADH and 18 μM riboflavins whose spectrum spans across all our spectral channels. The excitation and emission spectra of this calibration fluid was measured using the fluorimeter (FluroMax Plus-c; Horiba, Japan) and imaged on the hyperspectral microscope across all spectral channels. The calibration images were then used to correct the same spectrum measured with the hyperspectral system to be able to assign the unmixed fluorophores as well as to flatten the uneven illumination of fluorescence images generated by using our custom-made GUI software. Next, samples were manually segmented by using the DIC image taken concurrently with the hyperspectral images, creating a region of interest (ROI) in preparation for the unmixing process. For fibroblasts and individual blastomeres the ROI was drawn around the cell membrane that was clearly visible on the DIC image. For blastocysts, the ROI was manually drawn around the ICM by an experienced embryologist, avoiding the apical cells where TE might contaminate the signal.

Secondly, following image segmentation with generated region of interest (ROI) for each single channel image of each sample, the autofluorescence data were then subjected to feature analysis. The mathematical algorithms used here employed various transformations to capture various aspects of cell spectra and patterns in the cell images such as average intensity per cell of autofluorescent signals in each channel, the ratios of such average channel intensities for each pair pf channels (see Table III), various types of spatial, frequency, spectral, geometric, morphological and statistical information (Gosnell, *et al.*, 2016b); altogether approximately 33,000 cellular features were identified and used in the analysis. Next, depending on the selection of cell/embryo groups under consideration, indicative features which passed an ANOVA test (P < 0.005) were ranked according to their significance for classification into the specific groups, correlated features were removed, and the highest ranked uncorrelated features were selected as described previously (Gosnell, *et al.*, 2016a), yielding a different feature set for each different group comparison. A total of 10 features (see Table III for the list of features) with the highest ranking were selected for the classification of human cells (refer Figure 2) and 7 features were used for mouse individual blastomeres (refer Figure 4 and 5; D, and E) and blastocysts (refer Figure 6; D and E). These selected feature sets generated optimal separation between the groups under consideration. Further, these selected features for human cells, mouse blastomeres and blastocysts were projected onto a two-dimensional (2D) space created by linear discrimination analysis (LDA) (Jombart, *et al.*, 2010). This procedure maximises the distance between the two distinct groups of cells (euploid and aneuploid) across all cell types while minimizing the variance within each group (Jombart, *et al.*, 2010). The 2D space is spanned by two canonical variables that are linear combination of selected features described above (10 features for human cells; 7 features for mouse blastomeres and blastocysts) (Naganathan, *et al.*, 2008). Following this, a receiver operating characteristics (ROC) curve was generated and the area under the curve (AUC) was calculated to evaluate the accuracy of the separation between groups across cell types (Hand, *et al.*, 2001).

**Table III.** Number of features identified and used for analysis. Distinct features were selected according to experimental objectives and cell types listed as follows: 10 features were used to separate between euploid and aneuploid human fibroblast cells in Figure 2; 7 features were used to distinguish between euploid and aneuploid mouse blastomeres in Figure 4; euploid and aneuploid blastocysts in Figure 5D and 5E; and 1:1 and 1:3 chimeric blastocysts in Figure 6D and 6E.

| Figures in this manuscript | Features | Description of methodology used for analysis |
| --- | --- | --- |
| Figure 2 | 1 | Mean value of channel 26 divided by mean value of channel 58 |
|  | 2 | Mean value of channel 4 times by mean value of channel 10 |
|  | 3 | Contrast value of channel 63 |
|  | 4 | Mean value of channel 67 divided by mean value of channel 42 |
|  | 5 | Mean value of channel 50 divided by mean value of channel 9 |
|  | 6 | Mean value of channel 14 divided by mean value of channel 64 |
|  | 7 | Kurtosis value of second principal component image |
|  | 8 | Energy value of channel 63 |
|  | 9 | Mean value of second principal component image divided by mean value of channel 48 |
|  | 10 | Mean value of channel 19 divided by mean value of channel 52 |
| Figure 4 | 1 | Mean value of channel 55 divided by mean value of top 40 percent pixel values channel 37 |
|  | 2 | Mean value of first principal component image divided by mean value of top 40 percent pixel values of channel 46 |
|  | 3 | Mean value of first principal component image divided by mean value of channel 56 |
|  | 4 | Mean value of first principal component image divided by mean value of top 40 percent pixel values of channel 58 |
|  | 5 | Mean value of channel 67 |
|  | 6 | Mean value of first principal component image divided by mean value of channel 18 |
|  | 7 | Mean value of first principal component image divided by mean value of channel 32 |
| Figure 5D, 5E | 1 | Mean value of channel 47 divided by mean value of top 40 percent pixel values of channel 24 |
|  | 2 | Mean value channel of 47 divided by mean value of channel 5 |
|  | 3 | Variance value of channel 56 |
|  | 4 | Skewness value of channel 15 |
|  | 5 | Mean value of second principal component image divided by mean value of channel 18 |
|  | 6 | Skewness value of channel 38 |
|  | 7 | Kurtosis value of channel 28 |
| Figure 6D, 6E | 1 | Mean value of channel 28 |
|  | 2 | Mean value of channel 41 divided by mean value of top 40% channel 39 |
|  | 3 | Mean value channel 32 times by mean value of top 40% channel 40 |
|  | 4 | Skewness value of channel 60 |
|  | 5 | Mean value of second principal component image divided by mean value of third principal component image |
|  | 6 | Mean value of channel 8 divided by mean value of channel 36 |
|  | 7 | Mean value of channel 8 divided by mean value of top 40 percent pixel values channel 9 |

**Figure 1.**
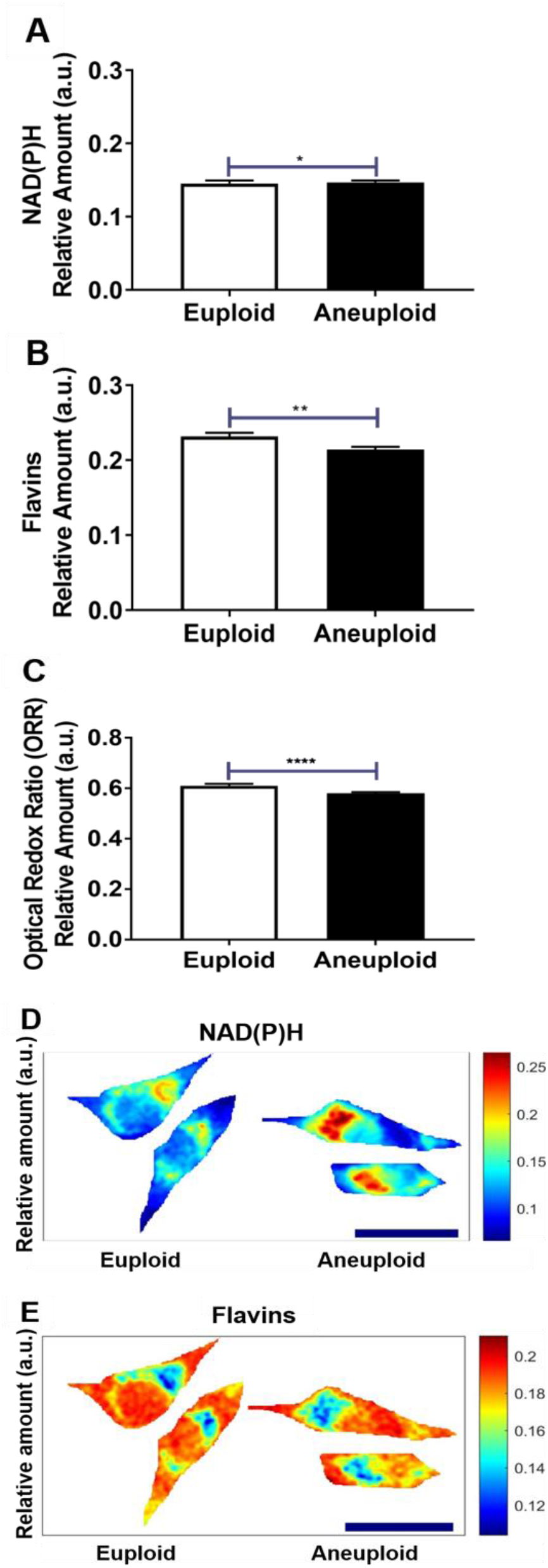
Metabolic activity is altered in aneuploid human primary fibroblasts. Primary human fibroblasts with known karyotypes (euploid [male and female] or aneuploid [triploid and trisomies: 13, 18, 21, XXX, and XXY]) were imaged using hyperspectral microscopy. Intracellular autofluorescence signals were subjected to unmixing for quantification of the relative abundance of NAD(P)H (A) and Flavins (B). The optical redox ratio (Flavins/ [NAD(P)H + Flavins])) was also calculated (C). Heat maps for euploid and aneuploid fibroblast cells were generated and show the localisation and abundance of fluorophores NAD(P)H and Flavins with representative cells shown in D and E, respectively. Scale bar = 20 μm. Data were analysed using Mann-Whitney test and presented as mean ± SEM. Four to six independent experimental replicates were performed; *n* = 467 for euploid cells; *n* = 969 for aneuploid cells. * *P* < 0.05, ** *P* < 0.01, **** *P* < 0.001.

**Figure 2.**
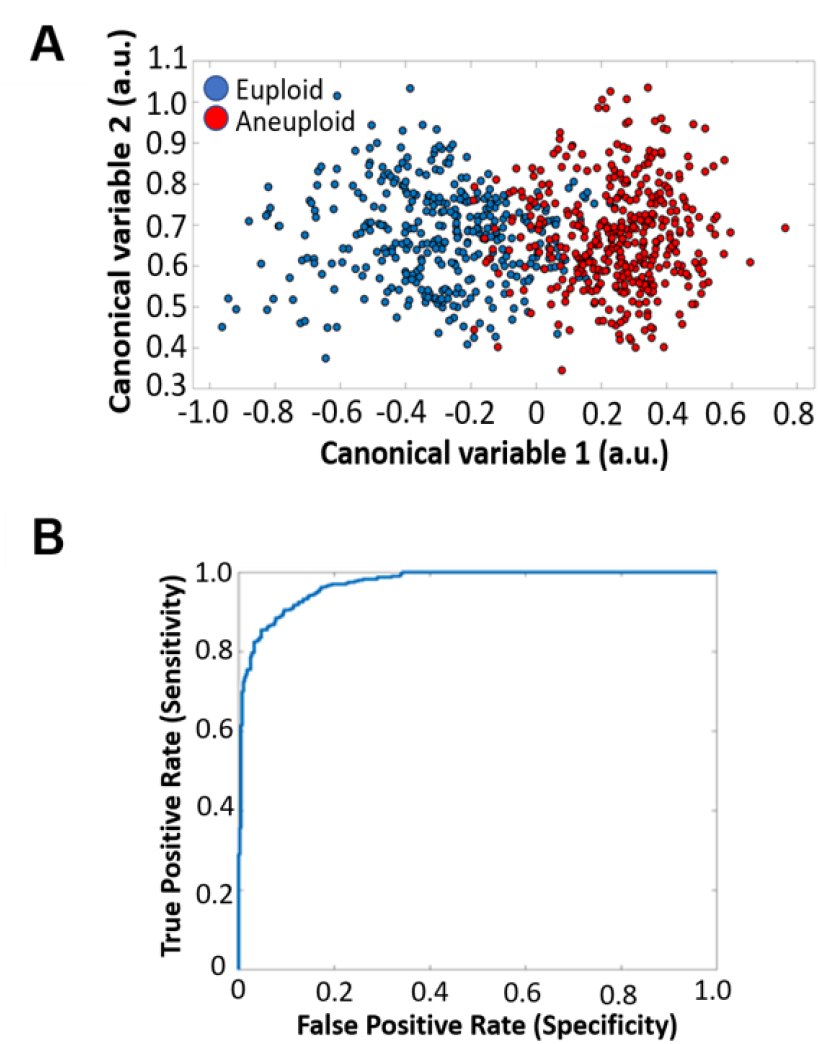
Mathematical algorithms separate euploid from aneuploid in primary human fibroblast cells. The autofluorescence data for each human fibroblast cell was subjected to feature analysis using a suite of mathematical algorithms as described in *materials and methods*. A cluster separation graph was generated using each cell (A, *blue*: male and female euploid cells; *red*: Triploid and Trisomies 13, 18, 21, XXX, XXY). Canonical values represent combinations of multiple features. (B) A receiver operating characteristic (ROC) curve was generated to determine the accuracy of separation between euploid and aneuploid cells (area under the curve (AUC) = 0.85). *n =* 364 for euploid cells; n= 400 for aneuploid cells.

**Figure 3:**
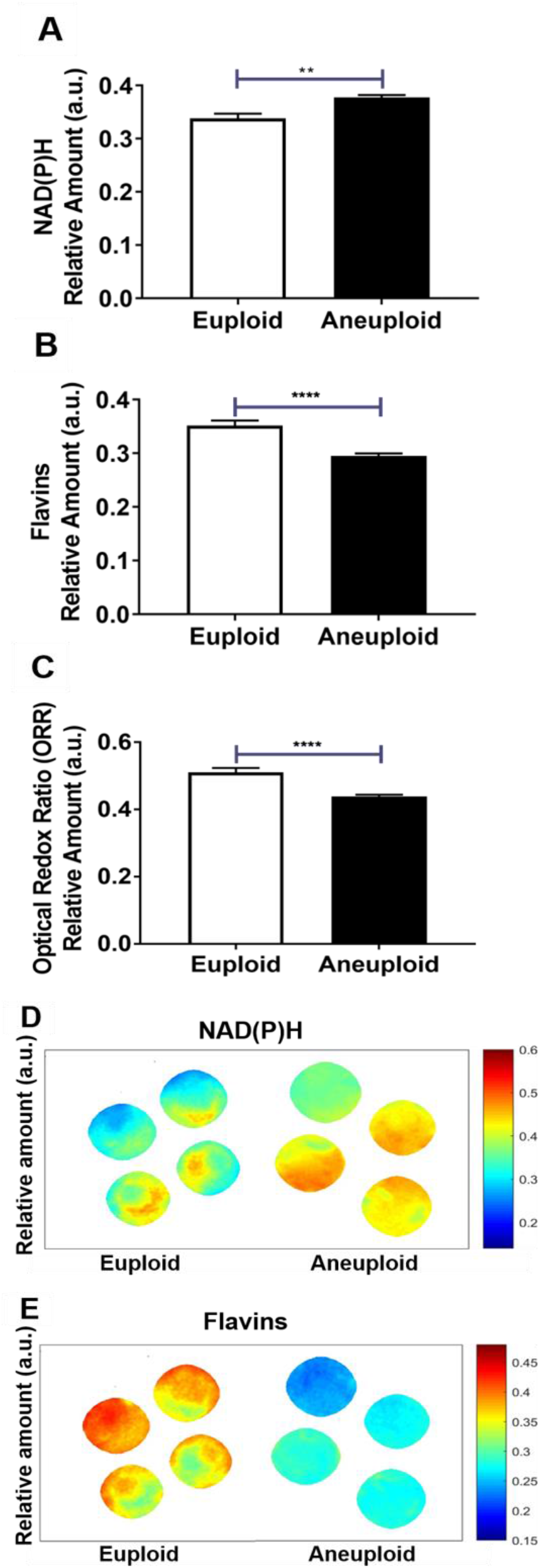
Aneuploidy results in an altered metabolism in mouse blastomeres. Aneuploidy was induced during the 4-to 8-cell division in the presence of reversine (0.5 uM). Eight-cell embryos were dissociated into individual blastomeres and imaged. Aneuploid blastomeres (reversine treated) were compared to euploid mouse blastomeres (control). Intracellular autofluorescence signals were subjected to unmixing for quantification of the relative abundance of NAD(P)H (A) and Flavins (B). The optical redox ratio (Flavins/ [NAD(P)H + Flavins]) was also calculated (C). Heat maps for euploid and aneuploid individual mouse blastomeres were generated and depict localisation and abundance of fluorophores NAD(P)H and Flavins with representative blastomeres shown in D and E, respectively. Scale bar = 20 μm. Data were analysed using Mann-Whitney test and presented as mean ± SEM. Five independent experimental r eplicates were pe rformed; *n* = 39 for e uploid; *n* = 44 fo r a neuploid blastom eres. * *P* < 0.05, **** *P* < 0.0 01.

**Figure 4:**
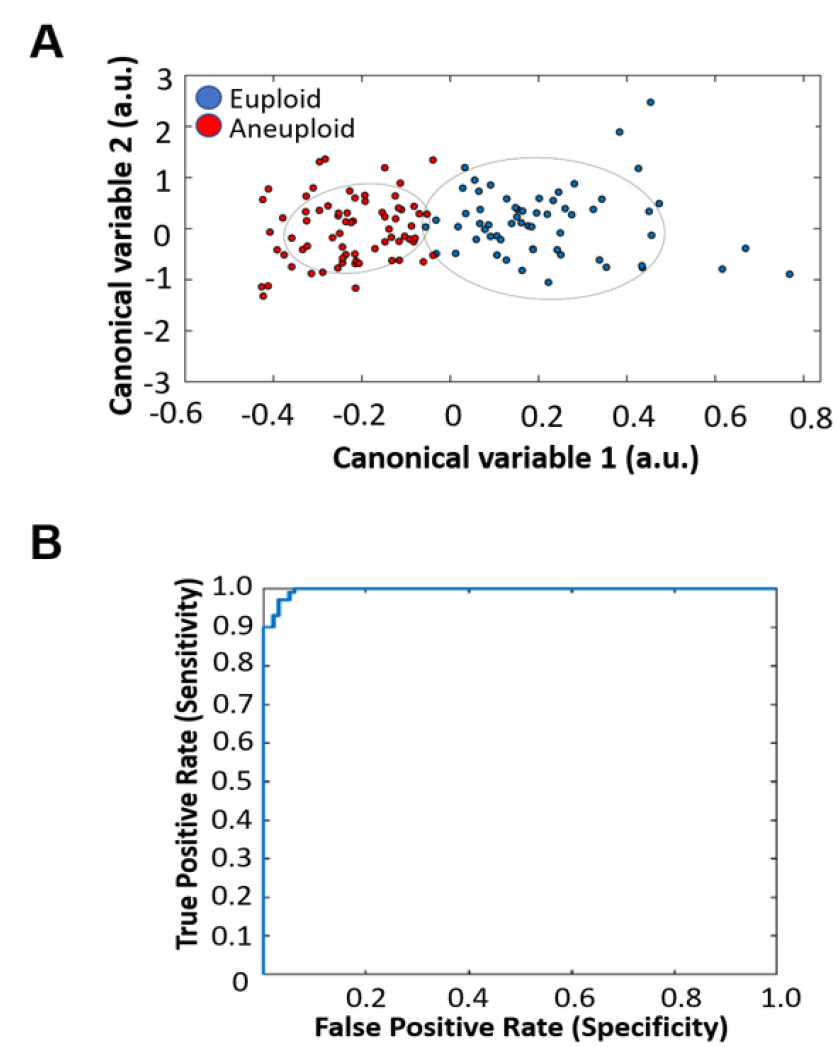
Mathematical algorithms separate euploid and aneuploid mouse blastomeres. The autofluorescence data for each mouse blastomere was subjected to feature analysis using mathematical algorithms as described in *materials and methods*. A cluster separation graph was generated using each cell (A, *blue*: euploid; *red*: aneuploid). Canonical values represent combinations of multiple features. (B) A receiver operating characteristic (ROC) curve was generated to determine the accuracy of separation between euploid and aneuploid cells mouse blastomeres (area under the curve (AUC) = 0.99). Five independent experimental replicates were p erformed; *n =* 61 euploid; *n* = 71 aneuploid blastomeres.

**Figure 5:**
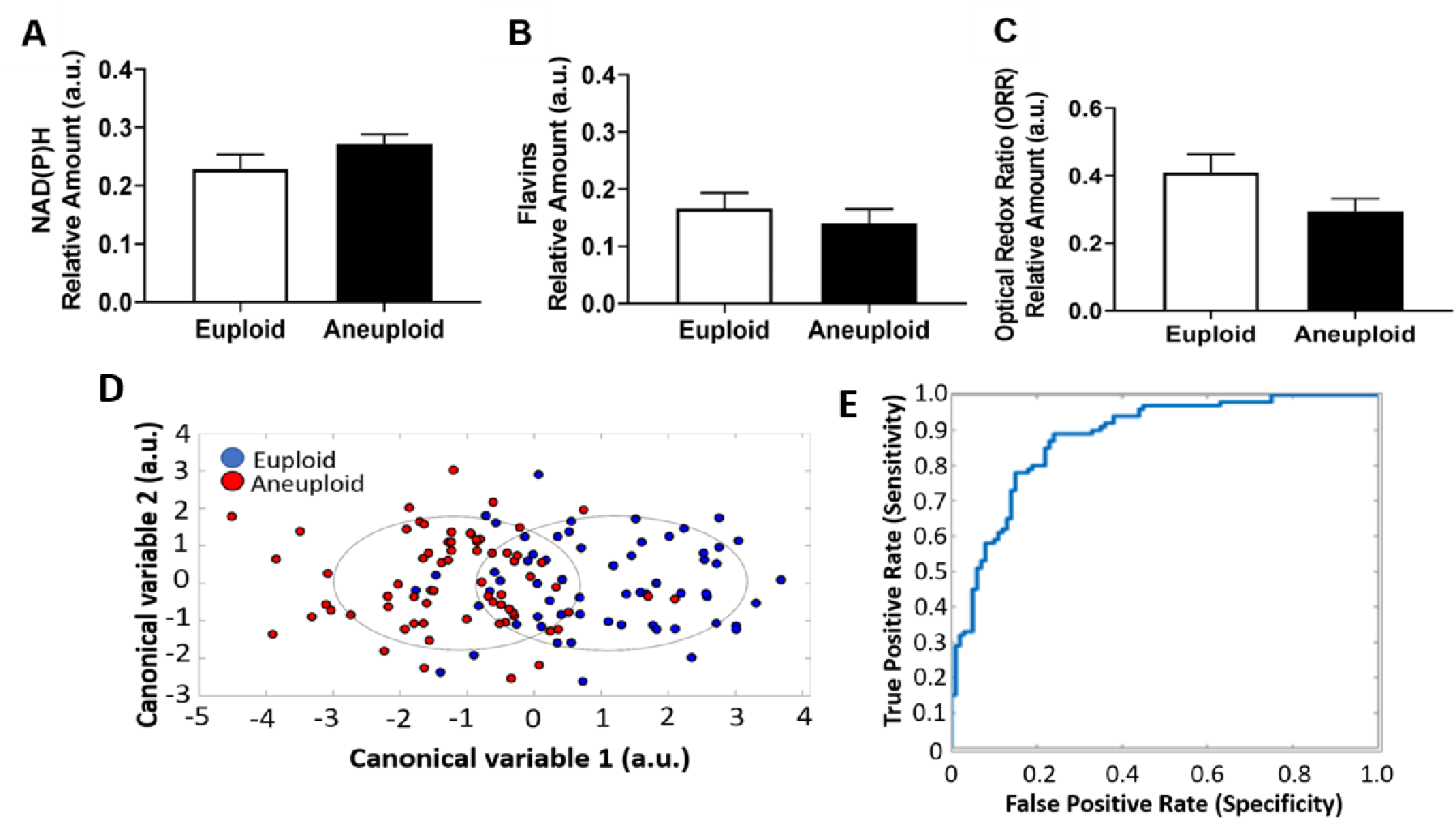
Euploid and aneuploid chimeric blastocysts show no difference in metabolic fluorophores but separation achieved using mathematical algorithms. Chimeric euploid and aneuploid blastocysts were generated as described in *materials and methods*. Chimeric blastocysts were subjected to hyperspectral imaging with field of view adjusted to focus on the inner cell mass (ICM). Intracellular autofluorescence signals were subjected to unmixing for quantification of the relative abundance of NAD(P)H (A) and Flavins (B). The optical redox ratio (Flavins/ [NAD(P)H + Flavins])) was also calculated (C). Autofluorescence spectra for each ICM was subjected to feature analysis using mathematical algorithms as described in *materials and methods*. A cluster separation graph was generated for euploid (*blue*) and aneuploid (*red*) chimeric blastocysts (D). Canonical values represent combinations of multiple features. (E) A receiver operating characteristic (ROC) curve was also generated to show separation between euploid and aneuploid ICM (area under the curve (AUC) = 0.87). Data in A-C are presented as mean ± SEM and were analysed by either unpaired Student’s t-test (A) or Mann-Whitney test (B and C). Eight independent experimental replicates were performed; *n =* 34 euploid; *n* = 39 aneuploid chimeric blastocysts

**Figure 6:**
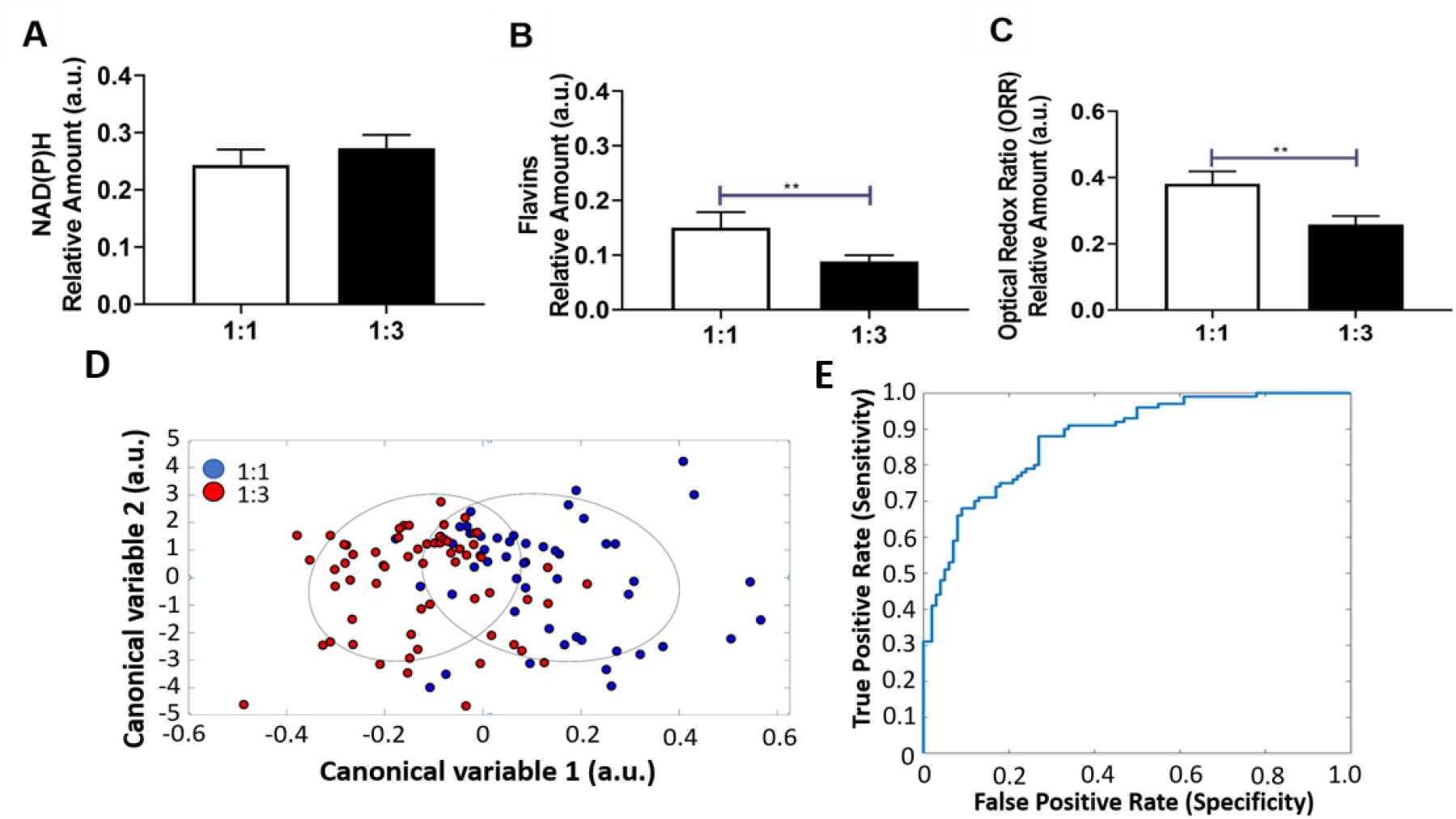
A higher ratio of aneuploid cells within the inner cell mass (ICM) of chimeric blastocysts leads to altered metabolism. Chimeric 1:1 and 1:3 (euploid:aneuploid) blastocysts were generated as described in *materials and methods* and subjected to hyperspectral imaging with field of view adjusted to focus on the ICM. Intracellular autofluorescence signals were unmixed to quantify of the relative a bundance of NAD(P) H (A) and Flavins (B). The optical redox ratio (Flavins/ [NAD(P)H + Flavins])) was also calculated (C). A cluster separation graph was generated for chimeric blastocysts to show separation between 1:1 (euploid:aneuploid, *blue*) and 1:3 (euploid:aneuploid, *red*) blastocysts (D). Canonical values represent combinations of multiple features. (E) A receiver operating characteristic (ROC) curve was generated to show separation between 1:1 and 1:3 blastocysts (area under the curve (AUC) = 0.83). Data in A-C are presented as mean ± SEM and were analysed by either unpaired Student’s t-test (A) or Mann-Whitney test (B and C). Eight independent experimental replicates were performed; *n =* 30 1:1 chimeric blastocysts; *n* = 37 1:3 chimeric blastocysts. ** *P* < 0.01.

Lastly, unmixing is the process where the spectral characteristic of the cells related to identified molecular fluorophores is extracted from the total autofluorescence signal. This is verified by comparison with known characteristics of the fluorophores at typical cellular concentrations and their relative abundance level (percentage of total fluorophore content) is calculated on a pixel by pixel basis. In this study, a linear mixing model (LMM) was adapted to perform the unmixing (Gosnell, *et al.*, 2016a, Mahbub, *et al.*, 2017). The model assumes that the fluorescent signal of each pixel is a linear combination of the endmember component spectra with their linear coefficients (weights) corresponding to the concentration of the molecules responsible for these component spectra. An unsupervised unmixing algorithm, Robust Dependent Component Analysis (RoDECA) (Mahbub, *et al.*, 2017) was then employed to detect the native fluorophores and calculate their relative abundance in the hyperspectral dataset across cell types. The accuracy of RoDECA in separating and quantifying the spectral signals of specific fluorophores from the complex noise in the cellular environment was established previously (Mahbub, *et al.*, 2019, Mahbub, *et al.*, 2017). Amongst the identified fluorophores across all cell types, metabolic co-factors - NAD(P)H and flavins were able to be identified (data not shown) and therefore were used for subsequent analysis for metabolic activity. The optical redox ratio was calculated and defined as the intensity of flavins divided by the sum of intensity of NAD(P)H and flavins which reflects the activity of the mitochondrial electron transport chain, and therefore of cellular metabolism (Kolenc, *et al.*, 2019, Yong, *et al.*, 2019)

### 2.9 Assessment of photodamage following hyperspectral imaging

Zona-enclosed embryos were generated by IVF and cultured as described above. Embryos at the morula stage of development were exposed to light in a hyperspectral microscope under identical conditions as during imaging at UoA (Table II). Non-imaged embryos were included and subjected to identical handling procedures as the imaged embryos (i.e. time out of the incubator). Both groups of embryos were cultured for a further 24 h when blastocyst development was scored, and embryos were assessed for DNA damage.

Imaged and non-imaged embryos were stained for γH2AX to quantify for DNA damage as previously described (Brown, *et al.*, 2018). Briefly, blastocysts were fixed in 4% paraformaldehyde (PFA) diluted in PBS for 30 min at room temperature. Following fixation, embryos were washed with PBV (PBS containing 0.3 mg/ml of polyvinyl alcohol) and permeabilized for 30 mins in 0.25 % v/v Triton-X100 in PBS. Embryos were then blocked with 10 % goat serum (Jackson Immuno, Philadelphia, Pennsylvania) diluted in PBV for 1 h. After blocking, embryos were incubated overnight at room temperature in the dark with anti-γH2AX primary antibody (Cell Signalling Technology, Danvers, MA) at 1:200 dilution with 10% goat serum in PBV. A negative control was included where embryos were incubated in the absence of the primary antibody. On the following day, embryos were washed 3 times in PBV before incubation for 2 h at room temperature in the dark with anti-rabbit Alexa Fluor 594-conjugated secondary antibody (Life Technologies, Carlsbad, CA) at 1:500 dilution with 10 % goat serum in PBV. Embryos were then counterstained with 3 mM of 4’,6-diamidino-2-phenylindole (DAPI). Lastly, embryos were washed 3 times in PBV and transferred onto a glass microscope slide with DAKO mounting medium (Dako Inc., Carpinteria, California) and enclosed with a coverslip using a spacer (ThermoFisher Scientific, Waltham, Massachusetts). Embryos were imaged on an Olympus FV3000 Confocal Laser Scanning microscope (Olympus, Tokyo, Japan). Embryos were excited at laser 405 nm (Emission Detection Wavelength: 430 nm to 470 nm) for DAPI and excited at laser 594 nm (Emission Detection Wavelength: 610 nm to 710 nm) for γH2AX. Image acquisition occurred at 60x magnification with immersion oil. A z-stack was captured for each embryo with maximum projection created and fluorescence intensity measured using Image J software (National Institute of Health) (Sutton-McDowall, *et al.*, 2015, Tan, *et al.*, 2016). Instrument settings were kept constant across all experimental replicates.

Embryo transfer experiments were conducted following hyperspectral imaging on the microscope at UNSW (Table I). Blastocyst embryos were transferred into the uterine horn of pseudo-pregnant Swiss female mice 2.5 days post-coitum with vasectomised males. On the day of transfer, blastocysts were warmed as described above. Embryo warming was performed 2 hours prior to embryo transfer to allow sufficient time for recuperation from potential heat-shock induced following warming. Post-warming survival of embryo was about 80-85 % for imaged and non-imaged groups. Embryo transfer was performed under anaesthesia with isoflurane. Eight morphologically normal, expanded blastocysts were transferred to each uterine horn (16 embryos transferred per mouse). Imaged and non-imaged embryos were transferred to different recipients. Number of pups from each female recipient was recorded on delivery and monitored for downstream offspring health. At post-natal day 21, offspring were weaned, weighed and assessed for gross facial abnormalities.

### 2.10 Statistical Analysis

All hyperspectral data analyses were carried out using Matlab (R2017b) software. All statistical analyses were carried out on GraphPad Prism Version 8 for Windows (GraphPad Holdings LLC, CA, USA). Data were subjected to normality testing using D’Agostino-Pearson Omnibus normality test prior to analysis. Data were either analysed by unpaired student t-test or Mann-Whitney test (due to data not following normal distribution), as indicated in the figure legends. Data are presented as mean ± standard error of mean (SEM). Statistical significance was set at P-value < 0.05.

## 3. Results

### Autofluorescence: imaging, analysis and cell discrimination of euploid and aneuploid primary human fibroblasts

We utilised primary human fibroblast cells with known karyotypes (euploid; triploid and trisomies: 13, 18, 21, XXX, and XXY) to determine whether hyperspectral microscopy can discriminate between euploid and aneuploid cells based on their autofluorescence spectra. Following hyperspectral imaging, linear unmixing was applied to determine the relative abundance of the metabolic co-factors NAD(P)H and flavins. Further, features were applied to the autofluorescence images to discriminate between euploid and aneuploid cells.

The abundance of NAD(P)H was found to be significantly higher in aneuploid compared to euploid fibroblast cells (Figure 1A, *P* < 0.05). In contrast, the abundance of flavins was significantly lower in aneuploid cells in comparison to euploid (Figure 1B, *P* < 0.01), which was reflected in a significantly lower optical redox ratio (Figure 1C, *P* < 0.0001). To further explore these differences, heat maps were generated to visualise the localisation and abundance of NAD(P)H and flavins within individual cells. As observed with the quantified data, the heat map shows a higher intensity of NAD(P)H in aneuploid cells compared to euploid (Figure 1D). A similar correlation was also observed for flavins (Figure 1E), with a higher abundance seen in euploid fibroblasts compared to aneuploid. Interestingly, there was heterogeneity in the location and abundance of fluorophores between cell type (aneuploid vs euploid) and within cells (Figure 1D and E).

Next, we extracted features from the hyperspectral data and conducted analysis which demonstrated that hyperspectral features can effectively discriminate between euploid and aneuploid human fibroblasts. Features identified are described in Table III. The cellular vectors of the selected and most discriminative features were plotted in a 2D discriminative space generated by LDA and span by two canonical variables which demonstrated clear clustering of euploid and aneuploid fibroblast cells (Figure 2A). To determine the accuracy of this separation, a ROC curve was generated and displayed a high accuracy in discriminating euploid and aneuploid fibroblast cells, with an AUC value of 0.85 (Figure 2B).

### Autofluorescence: imaging, analysis and cell discrimination of euploid and aneuploid primary mouse blastomeres

We next determined whether individual aneuploid blastomeres from mouse embryos showed a similar shift in the abundance of endogenous fluorophores and whether features analysis could separate euploid and aneuploid blastomeres. Reversine was used to induce aneuploidy during the 4-to 8-cell division of the embryo at which point individual blastomeres were dissociated and imaged. Compared to euploid blastomeres, the abundance of NAD(P)H in aneuploid blastomeres was significantly higher (Figure 3A, *P* < 0.01), while the abundance of flavins was significantly lower (Figure 3B, *P* < 0.0001). These changes were reflected in a significantly lower optical redox ratio in aneuploid blastomeres (Figure 3C; *P* < 0.0001) when compared to euploid blastomeres. Heat maps generated for the cellular distribution of NAD(P)H and flavins confirmed our findings in overall abundance for these metabolic co-factors between euploid and aneuploid individual blastomeres (Figure 3D and E).

The extraction of features from the hyperspectral data showed that the chosen algorithms could effectively discriminate between euploid and aneuploid mouse blastomeres. Features identified are described in Table III. As for human primary fibroblasts, cellular vectors of selected and discriminative features were plotted in a 2D space generated by LDA and span by two canonical variables showing a clear clustering of euploid and aneuploid blastomeres (Figure 4A). To determine the accuracy of this separation, a ROC curve was generated and displayed very high accuracy with an AUC value of 0.99 (Figure 4B).

### Autofluorescence: imaging, analysis and cell discrimination of mouse blastocysts with differing ratios of euploid:aneuploid cells

We next sought to determine whether the inner cell mass (ICM) of blastocyst embryos with differing proportions of aneuploid cells displayed differences in native fluorophores. We employed a model of euploid:aneuploid mosaicism (Bolton, *et al.*, 2016), where reversine was used to induce aneuploidy during the 4-to 8-cell division after which, individual blastomeres were dissociated and reaggregated with untreated (euploid) blastomeres to form the desired proportion of euploid:aneuploid (euploid; aneuploid; 1:1 or 1:3, respectively). Chimeric blastocyst embryos were imaged where the plane of interest, the ICM, was focused on. Unmixing of endogenous fluorophores showed no significant differences in the relative abundance of NAD(P)H (Figure 5A), flavins (Figure 5B) or the optical redox ratio (Figure 5C) between euploid and aneuploid chimeric blastocysts. Similarly, 1:1 and 1:3 chimeric blastocysts demonstrated no significant difference in NAD(P)H abundance level (Figure 6A). In contrast, the abundance of flavins was significantly lower in 1:3 chimeric blastocysts compared to 1:1 (Figure 6B, *P* < 0.001), which was reflected in a significantly lower optical redox ratio (Figure 6C, *P* < 0.001). The cellular vectors of the selected and most discriminative features (refer Table III for list of features) were plotted which demonstrated clear clustering, albeit with some degree of overlapping of euploid and aneuploid (Figure 5D) as well as 1:1 and 1:3 (Figure 6D). To determine the accuracy of this separation, a ROC curves were generated and displayed high accuracy in discriminating euploid and aneuploid chimeric blastocysts (Figure 5E; AUC value of 0.87) and 1:1 vs 1:3 (Figure 6E; AUC value of 0.88).

Due to the process of generating chimeric blastocysts, imaging and analysis occurred in the absence of a zona pellucida. We set out to confirm that the presence of the zona pellucida yielded similar results for the metabolic fluorophores, NAH(P)H and flavins. Zona pellucida enclosed blastocysts were imaged and compered to chimeric, zona free, blastocysts. Fluorophore unmixing showed no significant difference between zona pellucida-enclosed and zona pellucida-free embryos (data not shown).

### Safety of hyperspectral imaging

To assess whether photodamage had occurred in response to imaging, untreated morula stage mouse embryos were subjected to either hyperspectral imaging (*imaged*) or not imaged (*non-imaged*). Imaging of morula embryos did not affect their ability to develop to the blastocyst stage (Figure 7A) nor the level of DNA damage within the subsequent blastocyst embryo (Figure 7B) compared to non-imaged embryos.

**Figure 7:**
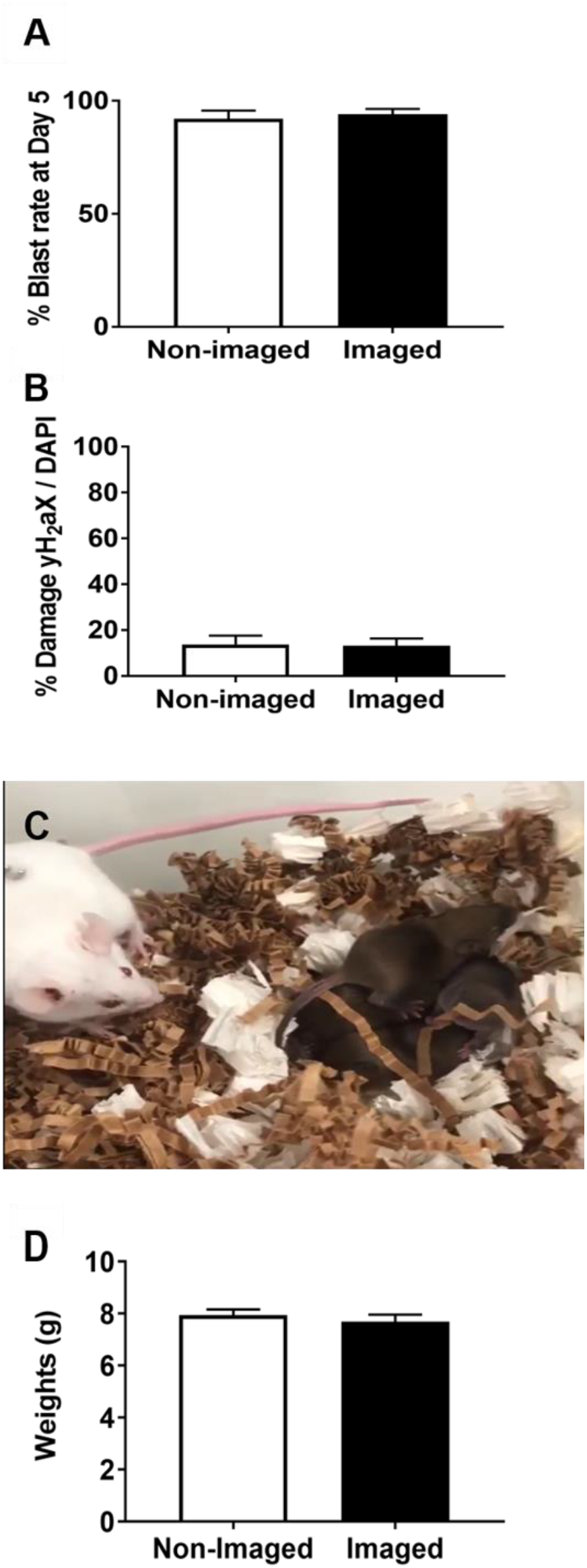
Hyperspectral imaging does not affect completion of preimplantation embryo development, the level of DNA damage within the embryo or postnatal outcomes following embryo transfer. Embryos at the morula stage were imaged using the hyperspectral icroscope and compared with non-imaged embryos for their capacity to reach the blastocyst stage of development (A) and the level of DNA damage within the blastocyst (B). Data were expressed as a percentage of γH2AX/DAPI (B). In a separate experiment, chimeric euploid blastocysts were generated and either imaged or not imaged using hyperspectral microscopy. Blastocysts were transferred to pseudo-pregnant females with imaged blastocysts resulting in the birth of live pups (C). Weights of pups at weaning (day 21) were assessed for each group (D). Data were analysed by Mann-Whitney test and presented as mean ± SEM, *n* = 50 embryos per group (A) and *n* = 11 embryos per group (B) representative of 5 experimental replicates. For D, *n* = 72 non-imaged and 88 imaged blastocysts transferred into 5 and 6 pseudopregnant recipients, respectively.

Next, we examined whether hyperspectral imaging impacted live birth rate or weight at weaning following transfer of imaged and non-imaged blastocysts to pseudopregnant mice. Imaged embryos resulted in the birth of live pups (Figure 7C). There was no significant difference in livebirth rate between non-imaged and imaged blastocysts (non-imaged: 47 full-term pups from 72 transferred embryos (65.3%); vs. imaged: 49 full-term pups from 80 transferred embryos (61.3%), data not shown). No gross facial deformities were noted across treatment groups. Finally, there was no significant difference in weight at weaning between pups derived from imaged and non-imaged embryos (Figure 7D).

## 4. Discussion

Clinical assessment of embryo aneuploidy remains a highly debatable topic due to the limitations and safety concerns of current approaches. Thus, there remains a need for an accurate, non-invasive clinical test for the detection and assessment of embryo aneuploidy. The present study showed that the metabolic activity of primary human fibroblasts, mouse blastomeres and the ICM of chimeric embryos was significantly altered due to the presence of aneuploidy. Further, we show that feature analysis of hyperspectral images was able to discriminate between euploid and aneuploid cells and embryos. Finally, we present data demonstrating that hyperspectral imaging poses no harm to embryo and offspring health using the assessments described herein.

Hyperspectral microscopy coupled with feature analysis demonstrated successful capture and identification of endogenous fluorophore abundance across different cell types with aneuploidy. In all cell types, aneuploidy was associated with an altered abundance of metabolic cofactors, which resulted in a lower optical redox ratio, and thus metabolic activity.

It is important to note the NAD(P)H fluorescence may be attributed to either NADH, cytosolic NADPH, or both, due to their near identical spectral properties (Galeotti, *et al.*, 1970, Rehman, *et al.*, 2017). Therefore, the results presented here could be ascribed to an increase in NADPH, which plays a key role in regulating oxidative stress, or NADH, which is essential for ATP synthesis via oxidative phosphorylation (reviewed in Ying, 2008). Elevated NAD(P)H may indicate increased oxidative stress due to an increase in oxidative phosphorylation, which generates reactive oxygen species (ROS) as a by-product. Supporting this, aneuploid mouse embryonic fibroblasts have been shown to contain higher levels of ROS than euploid cells (Li, *et al.*, 2010). Alternatively, the shift could also be due to a preferential shift to glycolysis, leading to increased NAD(P)H and decreased flavin abundance observed with aneuploidy. Indeed, cancer cells which have a high rate of aneuploidy are known for their dependence on glycolysis (Zhang, *et al.*, 2013) to satisfy the significant biosynthetic demands of uncontrolled proliferation, also known as the Warburg effect (Liberti, *et al.*, 2016). It is also interesting to note that in mouse embryos, a shift in metabolic pathway from oxidative phosphorylation to glycolysis occurs naturally during development to adapt to changes in micronutrient availability in the reproductive tract and energy demands of the embryo (Thompson, *et al.*, 2016). Therefore, it is tempting to postulate that aneuploidy may accelerate this process of change in metabolic pathway for mouse blastomeres and potentially chimeric blastocysts, leading to the observed altered metabolic profile. Collectively, our results align with the current understanding of how aneuploid cells display altered cellular metabolism compared to euploid cells (Williams, *et al.*, 2008, Zhu, *et al.*, 2018) and confirms the capability of label-free hyperspectral microscopy to discriminate cells based on cellular autofluorescence (Campbell, *et al.*, 2019, Gosnell, *et al.*, 2016b, Mahbub, *et al.*, 2019).

Furthermore, our results show the spatial localisation and abundance of endogenous fluorophores linked to metabolism. This provides further information on how fluorophores behave differently in aneuploid cells and that heterogeneity exists within and between cells. The different spatial profile of fluorophores in aneuploid cells observed in the current study may be associated with the altered localisation of chromosomes known to occur in aneuploid blastomeres (McKenzie, *et al.*, 2004). Overall, the results on spatial localisation of endogenous fluorophores demonstrates that aneuploid cells display an altered metabolic profile compared to euploid cells. To our knowledge, this is the first report of altered cell metabolism in mouse embryos in response to aneuploidy.

The safety of hyperspectral imaging was also assessed in the current study. This is of critical importance as existing literature shows that light exposure can be damaging to the embryo, negatively impacting preimplantation development and implantation, as well leading to increased levels of DNA damage within the embryo (Bognar, *et al.*, 2019, Oh, *et al.*, 2007, Takahashi, *et al.*, 1999). Our data showed that hyperspectral imaging did not affect blastocyst formation, level of DNA damage, or subsequent postnatal outcomes when compared to embryos that were not imaged. Collectively, these results demonstrate that hyperspectral imaging of the preimplantation embryo does not impact developmental competence, pregnancy outcome, and offspring health in a mouse model.

It is important to note that the present study utilised a mouse model to elucidate the impact of aneuploidy on metabolism and thus further interpretation or direct correlation to human embryos using these findings warrants caution. Furthermore, aneuploidy induced by reversine is random (Bolton, *et al.*, 2016), and may not reflect the types of aneuploidy observed clinically. However, we do not expect an effect of aneuploidy type on cell autofluorescence as variability in chromosome content yields a consistent transcriptional response in human cell lines compared to euploidy (Durrbaum, *et al.*, 2014). Accurate determination of the proportion of aneuploid cells within the ICM of individual blastocysts to correlate with altered metabolism and level of discrimination between euploid and aneuploid embryos is of interest. Finally, the predictive power of hyperspectral microscopy to detect differing proportions of aneuploid cells within the ICM vs TE and associate with subsequent developmental outcomes should be the basis of future study.

## 5. Conclusions

Hyperspectral imaging combined with feature analysis can accurately detect changes in metabolism associated with aneuploidy and discriminate between euploid and aneuploid human fibroblasts, mouse blastomeres, and blastocysts. No differences in developmental competence, DNA damage, and postnatal outcomes were observed in embryos subjected to hyperspectral imaging. Together, this work demonstrates that hyperspectral imaging of autofluorescence represents a new non-invasive approach to detect metabolic variance in embryos with aneuploidy. In the future, hyperspectral imaging of autofluorescence may lead to a more accurate and non-invasive assessment of the ploidy status of the divergent cell lineages of the blastocyst embryo.

## Author contributions

KRD conceived the idea for the study. TCYT, SBM, CAC, JMC, AH, SM, EG, and KRD were involved in experimental design and data analysis. TCYT, SBM, CAC, JMC and AH conducted experiments and performed data analysis. TCYT and KRD wrote the first draft of most of the manuscript, with specific sections written by SBM, AH, CAC, DJXC and EG. TCYT, SBM, AB and CAC generated the figures. All authors edited and approved the manuscript.

## Acknowledgements

The authors would like to thank Adelaide Microscopy at the University of Adelaide (Adelaide, South Australia, Australia) for their support throughout the duration of this study. The authors are also grateful to ART Lab solutions (Adelaide, South Australia, Australia) for their support in media use in this study. The authors also thank Dr Kishan Dholakia for his constructive comments during the preparation of the manuscript.

## References

Aparicio B, Cruz M, Meseguer M. Is morphokinetic analysis the answer? Reproductive BioMedicine Online 2013;27: 654–663.

Bertoldo MJ, Listijono DR, Ho WJ, Riepsamen AH, Goss DM, Richani D, Jin XL, Mahbub S, Campbell JM, Habibalahi A et al. NAD(+) Repletion Rescues Female Fertility during Reproductive Aging. Cell Rep 2020;30: 1670–1681 e1677.

Bognar Z, Csabai TJ, Pallinger E, Balassa T, Farkas N, Schmidt J, Gorgey E, Berta G, Szekeres-Bartho J, Bodis J. The effect of light exposure on the cleavage rate and implantation capacity of preimplantation murine embryos. J Reprod Immunol 2019;132: 21–28.

Bolton H, Graham SJL, Van der Aa N, Kumar P, Theunis K, Fernandez Gallardo E, Voet T, Zernicka-Goetz M. Mouse model of chromosome mosaicism reveals lineage-specific depletion of aneuploid cells and normal developmental potential. Nat Commun 2016;7: 11165.

Brown HM, Green ES, Tan TCY, Gonzalez MB, Rumbold AR, Hull ML, Norman RJ, Packer NH, Robertson SA, Thompson JG. Periconception onset diabetes is associated with embryopathy and fetal growth retardation, reproductive tract hyperglycosylation and impaired immune adaptation to pregnancy. Scientific Reports 2018;8.

Campbell JM, Habibalahi A, Mahbub S, Gosnell M, Anwer AG, Paton S, Gronthos S, Goldys E. Non-destructive, label free identification of cell cycle phase in cancer cells by multispectral microscopy of autofluorescence. BMC Cancer 2019;19: 1242.

Capalbo A, Rienzi L, Cimadomo D, Maggiulli R, Elliott T, Wright G, Nagy ZP, Ubaldi FM. Correlation between standard blastocyst morphology, euploidy and implantation: an observational study in two centers involving 956 screened blastocysts. Hum Reprod 2014;29: 1173–1181.

Dumollard R. Sperm-triggered [Ca2+] oscillations and Ca2+ homeostasis in the mouse egg have an absolute requirement for mitochondrial ATP production. Development 2004;131: 3057–3067.

Durrbaum M, Kuznetsova AY, Passerini V, Stingele S, Stoehr G, Storchova Z. Unique features of the transcriptional response to model aneuploidy in human cells. BMC Genomics 2014;15: 139.

Feichtinger M, Vaccari E, Carli L, Wallner E, Madel U, Figl K, Palini S, Feichtinger W. Non-invasive preimplantation genetic screening using array comparative genomic hybridization on spent culture media: a proof-of-concept pilot study. Reprod Biomed Online 2017;34: 583–589.

Fragouli E, Alfarawati S, Daphnis DD, Goodall NN, Mania A, Griffiths T, Gordon A, Wells D. Cytogenetic analysis of human blastocysts with the use of FISH, CGH and aCGH: scientific data and technical evaluation. Hum Reprod 2011;26: 480–490.

Fragouli E, Spath K, Alfarawati S, Kaper F, Craig A, Michel CE, Kokocinski F, Cohen J, Munne S, Wells D. Altered levels of mitochondrial DNA are associated with female age, aneuploidy, and provide an independent measure of embryonic implantation potential. PLoS Genet 2015;11: e1005241.

Galeotti T, van Rossum GD, Mayer DH, Chance B. On the fluorescence of NAD(P)H in whole-cell preparations of tumours and normal tissues. Eur J Biochem 1970;17: 485–496.

Gleicher N, Barad DH. Not even noninvasive cell-free DNA can rescue preimplantation genetic testing. Proc Natl Acad Sci U S A 2019;116: 21976–21977.

Gleicher N, Metzger J, Croft G, Kushnir VA, Albertini DF, Barad DH. A single trophectoderm biopsy at blastocyst stage is mathematically unable to determine embryo ploidy accurately enough for clinical use. Reprod Biol Endocrinol 2017;15: 33.

Gosnell ME, Anwer AG, Cassano JC, Sue CM, Goldys EM. Functional hyperspectral imaging captures subtle details of cell metabolism in olfactory neurosphere cells, disease-specific models of neurodegenerative disorders. Biochim Biophys Acta 2016a;1863: 56–63.

Gosnell ME, Anwer AG, Mahbub SB, Menon Perinchery S, Inglis DW, Adhikary PP, Jazayeri JA, Cahill MA, Saad S, Pollock CA et al. Quantitative non-invasive cell characterisation and discrimination based on multispectral autofluorescence features. Sci Rep 2016b;6: 23453.

Habibalahi A, Bala C, Allende A, Anwer AG, Goldys EM. Novel automated non invasive detection of ocular surface squamous neoplasia using multispectral autofluorescence imaging. Ocul Surf 2019;17: 540–550.

Hand DJ, Till RJ. A Simple Generalisation of the Area Under the ROC Curve for Multiple Class Classification Problems. Machine Learning 2001;45: 171–186.

Huang L, Bogale B, Tang Y, Lu S, Xie XS, Racowsky C. Noninvasive preimplantation genetic testing for aneuploidy in spent medium may be more reliable than trophectoderm biopsy. Proc Natl Acad Sci U S A 2019;116: 14105–14112.

Johnson DS, Cinnioglu C, Ross R, Filby A, Gemelos G, Hill M, Ryan A, Smotrich D, Rabinowitz M, Murray MJ. Comprehensive analysis of karyotypic mosaicism between trophectoderm and inner cell mass. Mol Hum Reprod 2010;16: 944–949.

Jombart T, Devillard S, Balloux F. Discriminant analysis of principal components: a new method for the analysis of genetically structured populations. BMC Genetics 2010;11: 94.

Kolenc OI, Quinn KP. Evaluating Cell Metabolism Through Autofluorescence Imaging of NAD(P)H and FAD. Antioxidants & Redox Signaling 2019;30: 875–889.

Kuznyetsov V, Madjunkova S, Abramov R, Antes R, Ibarrientos Z, Motamedi G, Zaman A, Kuznyetsova I, Librach CL. Minimally Invasive Cell-Free Human Embryo Aneuploidy Testing (miPGT-A) Utilizing Combined Spent Embryo Culture Medium and Blastocoel Fluid-Towards Development of a Clinical Assay. Sci Rep 2020;10: 7244.

Li M, Fang X, Baker DJ, Guo L, Gao X, Wei Z, Han S, van Deursen JM, Zhang P. The ATM-p53 pathway suppresses aneuploidy-induced tumorigenesis. Proceedings of the National Academy of Sciences of the United States of America 2010;107: 14188–14193.

Liberti MV, Locasale JW. The Warburg Effect: How Does it Benefit Cancer Cells? Trends in Biochemical Sciences 2016;41: 211–218.

Mahbub SB, Guller A, Campbell JM, Anwer AG, Gosnell ME, Vesey G, Goldys EM. Non-Invasive Monitoring of Functional State of Articular Cartilage Tissue with Label-Free Unsupervised Hyperspectral Imaging. Scientific Reports 2019;9.

Mahbub SB, Ploschner M, Gosnell ME, Anwer AG, Goldys EM. Statistically strong label-free quantitative identification of native fluorophores in a biological sample. Sci Rep 2017;7: 15792.

Mastenbroek S, Twisk M, van der Veen F, Repping S. Preimplantation genetic screening: a systematic review and meta-analysis of RCTs. Hum Reprod Update 2011;17: 454–466.

McKenzie LJ, Carson SA, Marcelli S, Rooney E, Cisneros P, Torskey S, Buster J, Simpson JL, Bischoff FZ. Nuclear chromosomal localization in human preimplantation embryos: correlation with aneuploidy and embryo morphology. Hum Reprod 2004;19: 2231–2237.

Naganathan GK, Grimes LM, Subbiah J, Calkins CR, Samal A, Meyer GE. Visible/near-infrared hyperspectral imaging for beef tenderness prediction. Computers and Electronics in Agriculture 2008;64: 225–233.

Newman DL, Gregory SL. Co-Operation between Aneuploidy and Metabolic Changes in Driving Tumorigenesis. International journal of molecular sciences 2019;20: 4611.

Northrop LE, Treff NR, Levy B, Scott RT, Jr. SNP microarray-based 24 chromosome aneuploidy screening demonstrates that cleavage-stage FISH poorly predicts aneuploidy in embryos that develop to morphologically normal blastocysts. Mol Hum Reprod 2010;16: 590–600.

Oh SJ, Gong SP, Lee ST, Lee EJ, Lim JM. Light intensity and wavelength during embryo manipulation are important factors for maintaining viability of preimplantation embryos in vitro. Fertil Steril 2007;88: 1150–1157.

Picton HM, Elder K, Houghton FD, Hawkhead JA, Rutherford AJ, Hogg JE, Leese HJ, Harris SE. Association between amino acid turnover and chromosome aneuploidy during human preimplantation embryo development in vitro. Mol Hum Reprod 2010;16: 557–569.

Rehman AU, Anwer AG, Gosnell ME, Mahbub SB, Liu G, Goldys EM. Fluorescence quenching of free and bound NADH in HeLa cells determined by hyperspectral imaging and unmixing of cell autofluorescence. Biomedical Optics Express 2017;8: 1488.

Rubio C, Rienzi L, Navarro-Sanchez L, Cimadomo D, Garcia-Pascual CM, Albricci L, Soscia D, Valbuena D, Capalbo A, Ubaldi F et al. Embryonic cell-free DNA versus trophectoderm biopsy for aneuploidy testing: concordance rate and clinical implications. Fertil Steril 2019;112: 510–519.

Santaguida S, Tighe A, D’Alise AM, Taylor SS, Musacchio A. Dissecting the role of MPS1 in chromosome biorientation and the spindle checkpoint through the small molecule inhibitor reversine. J Cell Biol 2010;190: 73–87.

Santos Monteiro CA, Chow DJX, Leal GR, Tan TC, Reis Ferreira AM, Thompson JG, Dunning KR. Optical imaging of cleavage stage bovine embryos using hyperspectral and confocal approaches reveals metabolic differences between on-time and fast-developing embryos. Theriogenology 2020;159: 60–68.

Sheltzer JM. A transcriptional and metabolic signature of primary aneuploidy is present in chromosomally unstable cancer cells and informs clinical prognosis. Cancer Res 2013;73: 6401–6412.

Sheltzer JM, Blank HM, Pfau SJ, Tange Y, George BM, Humpton TJ, Brito IL, Hiraoka Y, Niwa O, Amon A. Aneuploidy drives genomic instability in yeast. Science (New York, NY) 2011;333: 1026–1030.

Sugimura S, Ritter LJ, Sutton-McDowall ML, Mottershead DG, Thompson JG, Gilchrist RB. Amphiregulin co-operates with bone morphogenetic protein 15 to increase bovine oocyte developmental competence: effects on gap junction-mediated metabolite supply. MHR: Basic science of reproductive medicine 2014;20: 499–513.

Sutton-McDowall ML, Gosnell M, Anwer AG, White M, Purdey M, Abell AD, Goldys EM, Thompson JG. Hyperspectral microscopy can detect metabolic heterogeneity within bovine post-compaction embryos incubated under two oxygen concentrations (7% versus 20%). Hum Reprod 2017;32: 2016–2025.

Sutton-McDowall ML, Purdey M, Brown HM, Abell AD, Mottershead DG, Cetica PD, Dalvit GC, Goldys EM, Gilchrist RB, Gardner DK et al. Redox and anti-oxidant state within cattle oocytes following in vitro maturation with bone morphogenetic protein 15 and follicle stimulating hormone. Molecular Reproduction and Development 2015;82: 281–294.

Sutton-McDowall ML, Wu LLY, Purdey M, Abell AD, Goldys EM, MacMillan KL, Thompson JG, Robker RL. Nonesterified Fatty Acid-Induced Endoplasmic Reticulum Stress in Cattle Cumulus Oocyte Complexes Alters Cell Metabolism and Developmental Competence1. Biology of Reproduction 2016;94.

Takahashi M, Saka N, Takahashi H, Kanai Y, Schultz RM, Okano A. Assessment of DNA damage in individual hamster embryos by comet assay. Mol Reprod Dev 1999;54: 1–7.

Tan TCY, Ritter LJ, Whitty A, Fernandez RC, Moran LJ, Robertson SA, Thompson JG, Brown HM. Gray level Co-occurrence Matrices (GLCM) to assess microstructural and textural changes in pre-implantation embryos. Molecular Reproduction and Development 2016;83: 701–713.

Taylor TH, Griffin DK, Katz SL, Crain JL, Johnson L, Gitlin S. Technique to ‘Map’ Chromosomal Mosaicism at the Blastocyst Stage. Cytogenetic and Genome Research 2016;149: 262–266.

Thompson JG, Brown HM, Sutton-McDowall ML. Measuring embryo metabolism to predict embryo quality. Reproduction, Fertility and Development 2016;28: 41–50.

Torres EM, Sokolsky T, Tucker CM, Chan LY, Boselli M, Dunham MJ, Amon A. Effects of aneuploidy on cellular physiology and cell division in haploid yeast. Science 2007;317: 916–924.

Vera-Rodriguez M, Diez-Juan A, Jimenez-Almazan J, Martinez S, Navarro R, Peinado V, Mercader A, Meseguer M, Blesa D, Moreno I et al. Origin and composition of cell-free DNA in spent medium from human embryo culture during preimplantation development. Hum Reprod 2018;33: 745–756.

Vera-Rodriguez M, Rubio C. Assessing the true incidence of mosaicism in preimplantation embryos. Fertil Steril 2017;107: 1107–1112.

Warburg O. On the origin of cancer cells. Science 1956;123: 309–314.

Williams BR, Prabhu VR, Hunter KE, Glazier CM, Whittaker CA, Housman DE, Amon A. Aneuploidy affects proliferation and spontaneous immortalization in mammalian cells. Science 2008;322: 703–709.

Xu J, Fang R, Chen L, Chen D, Xiao JP, Yang W, Wang H, Song X, Ma T, Bo S et al. Noninvasive chromosome screening of human embryos by genome sequencing of embryo culture medium for in vitro fertilization. Proc Natl Acad Sci U S A 2016;113: 11907–11912.

Ying W. NAD+/NADH and NADP+/NADPH in Cellular Functions and Cell Death: Regulation and Biological Consequences. Antioxidants & Redox Signaling 2008;10: 179–206.

Yong D, Abdul Rahim AA, Thwin CS, Chen S, Zhai W, Win Naing M. Autofluorescence spectroscopy in redox monitoring across cell confluencies. PLOS ONE 2019;14: e0226757.

Zhang Y, Yang J-M. Altered energy metabolism in cancer. Cancer Biology & Therapy 2013;14: 81–89.

Zhu J, Tsai HJ, Gordon MR, Li R. Cellular Stress Associated with Aneuploidy. Developmental Cell 2018;44: 420–431.

